# Development of Modular Expression and Genome Editing Across Phylogenetically Distinct Diazotrophs

**DOI:** 10.1101/2024.05.22.595406

**Authors:** Shawn Kulakowski, Alex Rivier, Rita Kuo, Sonya Mengel, Thomas Eng

## Abstract

Diazotrophic bacteria can reduce atmospheric nitrogen into ammonia enabling bioavailability of the essential element. Many diazotrophs associate closely with plant roots increasing nitrogen availability, acting as plant growth promoters. These associations have the potential to reduce the need for costly synthetic fertilizers if they could be engineered for agricultural applications. However, despite the importance of diazotrophic bacteria, genetic tools are poorly developed in a limited number of species, in turn narrowing the crops and root microbiomes that can be targeted. Here we report optimized protocols and plasmids to manipulate phylogenetically diverse diazotrophs with the goal of enabling synthetic biology and genetic engineering. Three broad-host-range plasmids can be used across multiple diazotrophs, with the identification of one specific plasmid (containing origin of replication RK2 and a kanamycin resistance marker) showing the highest degree of compatibility across bacteria tested. We then demonstrated modular expression testing seven promoters and eleven ribosomal binding sites using proxy fluorescent proteins. Finally, we tested four small molecule inducible systems to report expression in three diazotrophs and demonstrate genome editing in *Klebsiella michiganensis* M5al.

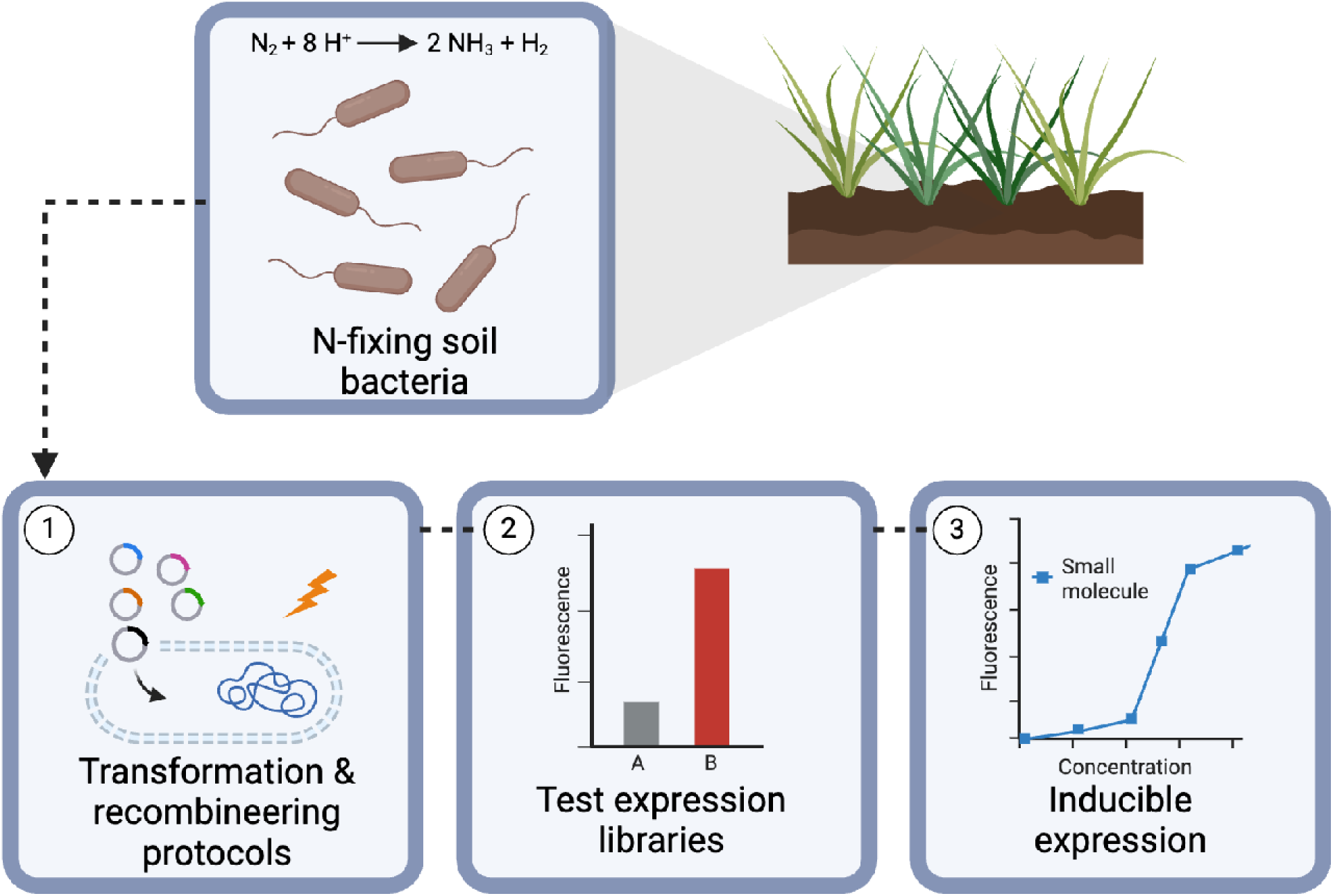

## Introduction

Nitrogen is an established key nutrient for the development of plants and has a significant effect on crop yield <u>(Lawlor et al. 2001)</u>. As a result, synthetic nitrogen sources are often added to improve crop performance. However, these synthetic fertilizers are generated using the energy intensive Haber-Bosch process which is estimated to use ∼2% of the world’s energy output <u>(Erisman et al. 2008)</u>. Synthetic fertilizers are also linked to negative environmental factors such as soil degradation and greenhouse gas emissions <u>(Kongshaug 1998; Sutton et al. 2011; Tripathi et al. 2020)</u>. Consequently, alternative nitrogen sources have become economically and environmentally attractive. Nitrogen-fixing bacteria, referred to as diazotrophs, possess the ability to perform biological nitrogen fixation (BNF), a potential route to lessen dependence on synthetic fertilizers. Diazotrophs can associate with crops and supply usable nitrogen <u>(Fernandes and Rossiello 1995; Robertson and Vitousek 2009)</u>. Research in diazotrophs has aimed to increase the nitrogen supplied to crops and also to further understand mechanisms that lead to and sustain plant-diazotroph associations or microbe-microbe interactions in the rhizosphere to improve overall nitrogen release productivity <u>(Pankievicz et al. 2019; Chakraborty et al. 2023; Knights et al. 2021)</u>. Despite the potential role for engineering diazotrophs for improved BNF, only a limited set of tools have been developed in a few diazotroph species <u>(Venkataraman et al. 2023)</u>. Since diazotrophs are known to be highly diverse, exist in variable ecological niches, and vary in plant association, it is desirable to have genetic tools that can be applied to a wide range of diazotrophs <u>(Pankievicz et al. 2019)</u>.

Our objective in this study was to build synthetic biology tools in a variety of diazotrophs across the proteobacterial clade to onboard potential model strains. We present here the development of plasmid-based expression systems in five diazotrophic bacterial species: *Klebsiella michiganensis* M5al (formerly referred to *Klebsiella oxytoca* M5al and *Klebsiella pneumonia* m5al) <u>(Mahl et al. 1965)</u>, *Azospirillum brasilense* Sp245 <u>(Baldani et al. 1983)</u>*, Herbaspirillum seropedicae* SmR1 <u>(Pedrosa et al. 2011)</u>*, Azorhizobium caulinodans* ORS 571 <u>(Lee et al. 2008; Dreyfus and Dommergues 1981)</u>, and *Rhizobium leguminosarum* 3Hoq18 <u>(Ramírez-Bahena et al. 2008)</u>. The relative phylogenetic relationships of our selected diazotrophs along with native host and their BNF modality are summarized in **Figure 1**. Two of the selected diazotrophs (*R. leguminosarum,* and *A. caulinodans*) are of the order Rhizobiales and symbiotically form root nodules in legumes where high rates of BNF occur <u>(Döbereiner 1997)</u>. *R. leguminosarum* was originally isolated from pea nodules and has been shown to be as effective as fertilizer on sweet pea inoculants <u>(Bedrous and Owais 1982)</u>. *A. caulinodans* is unusual in its ability to reduce nitrogen not only in tropical sesbania root nodules but also as a free-living soil microbe <u>(Lee et al. 2008)</u>. Additionally, *A. caulinodans* has the ability to colonize wheat roots, increasing dry weight and nitrogen content, which makes it suitable for studying both legume and grass interactions <u>(Sabry et al. 1997)</u>. More recently, *A. caulinodans* has been used to express nitrogenase systems under the control of synthetic bacterial signals engineered into *Medicago truncatula* and barley <u>(Geddes et al. 2019)</u>.

**Figure 1.**
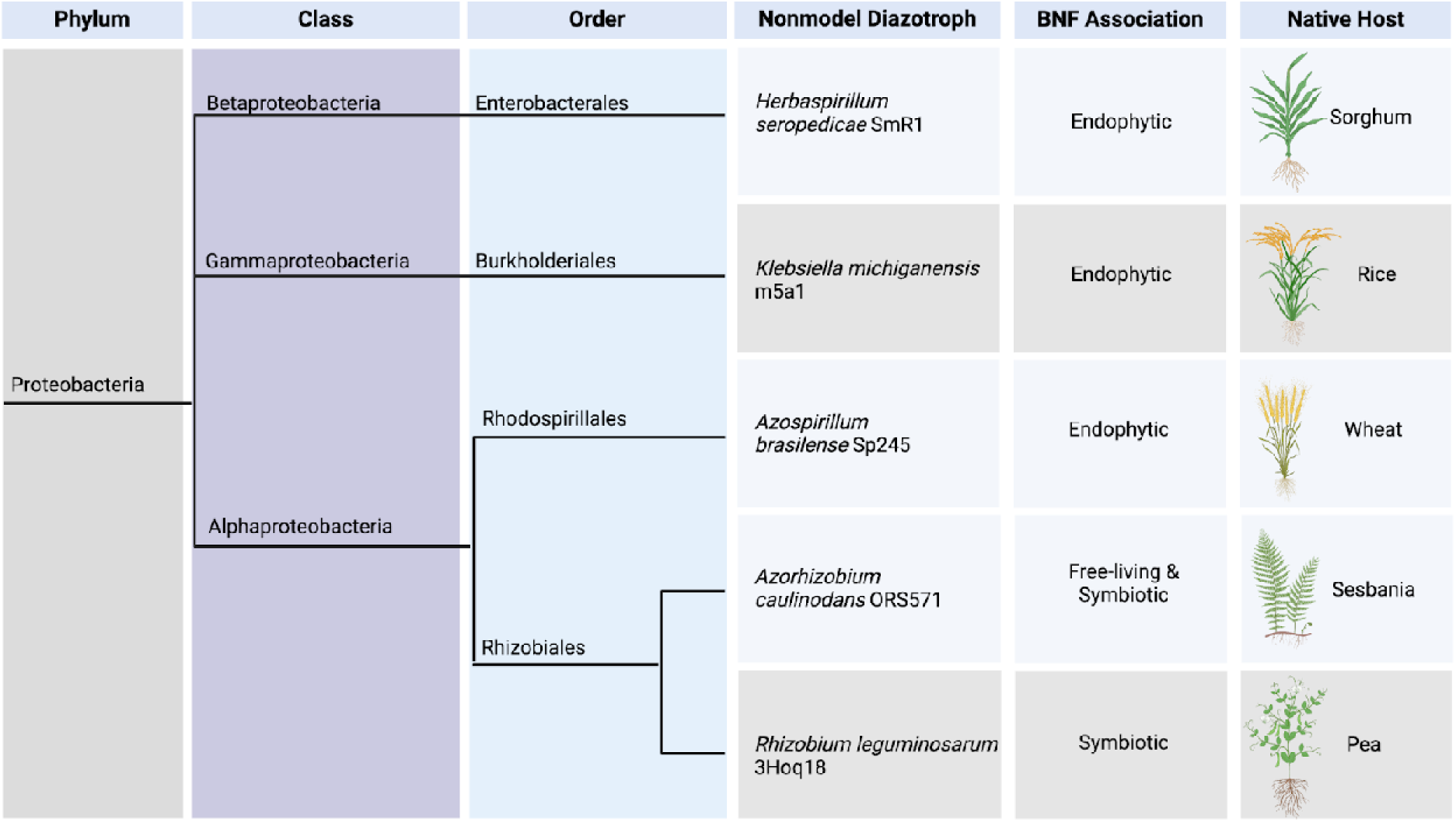
Graphical representation of phylogenetic distribution. From left to right: the relativ phylum, class, and order of each diazotroph (phylogenetic tree not to scale) followed by strain, nitrogen fixation modality (BNF association) as either endophytic (associated with roots), free-living (independent of plants), or symbiotic (root nodule symbiosis), and the native host from which the strain was isolated.

The other three diazotrophs (*K. michiganensis, A. brasilense,* and *H. seropedicae*) are phylogenetically dispersed across alpha-, gamma-, and beta-proteobacteria and are known to form endophytic relationships with a variety of grasses; colonizing in and around root structures <u>(Yu et al. 2018; Orlandini et al. 2014; Baldani et al. 1986)</u>. *K. michiganensis* strain m5al has long been a model for studying the molecular genetics of nitrogen fixation <u>(Parejko and Wilson 1970; Lin et al. 1993; Dixon and Postgate 1971)</u>. *K. michiganensis* has also been shown to produce cell wall-degrading compounds and colonize rice seedling roots <u>(Yu et al. 2018)</u>. *A. brasilense* natively associates with wheat and other grasses and has been shown to increase vegetative development of grape vines <u>(Bartolini et al. 2017)</u>. Lastly, *H. seropedicae,* an endophyte of tropical grasses, is reported to increase crop biomass and a range of secretion systems suggesting high interaction with plants <u>(Alberton et al. 2013; Pedrosa et al. 2011; Silveira Alves et al. 2019)</u> Overall, endophytes make promising model organisms given their ability to associate in close proximity with crops to supply nitrogen with an understanding of how sustained diazotroph-plant associations can be maintained.

Though all of the selected diazotrophs have been subject to experimentation they lack the support of tested genetic parts for stable, modular expression of exogenous DNA – an important yet time-consuming step for model strain onboarding due to the unbound experimental space needed to optimize parameters for growth and transformation. We focused on the transformation of broad-host plasmids using fluorescent markers as a proxy for heterologous gene expression. To modulate transcription and translation we queried alternate promoter sequences and ribosomal binding sites. We show inducible gene expression with signaling molecules known to be present in root exudates. Finally, we demonstrate genome editing in *K. michiganensis* with a newly constructed recombineering system. Our results enable the facile expression and manipulation of five diverse diazotrophs. Our tools and methods can be leveraged for the rapid assessment of genetic components integral to sustained diazotroph-plant interactions and nitrogen release of diazotrophs and ultimately, enable large scale commercial adoption of sustainable nitrogen sources.

## Results & Discussion

### Baseline Characterization of Diazotrophs for Lab Cultivation and Strain Engineering

We examined the literature and existing sequence databases to identify diazotrophs isolated from varied climates since the earliest reports in the 1940s <u>(Wilson and Burris 1947)</u>. From this information we were able to down select several diazotrophs on the basis of their availability in public repositories, existence of high-quality genomic sequencing information, and reports indicating cultivation under primarily aerobic laboratory conditions. Strain origins and accession numbers for American Type Culture Collection (ATCC), German Collection of Microorganisms and Cell Cultures (DSMZ), and National Center for Biotechnology Information (NCBI) accession numbers are summarized in **Supplemental Table 1**. Likely due to differences in strain collection standards over the past decades, exact geographical and climate isolation information is missing from several isolates. Additionally, laboratory cultivation of diazotrophs is often poorly characterized in the accompanying repository metadata, requiring further experimentation to onboard strains. More recent strain sequencing information indicates relatively large bacterial genome sizes (∼6Mb) but all have high quality scores as calculated by NCBI Prokaryotic Genome Annotation Pipeline (PGAP) <u>(Parks et al. 2015)</u> and fully assembled genomes. The GC content of the strains ranges from 56% to 68.5% in *K. michiganensis* and *A. brasilense*, respectively. All strains contain one chromosome but *A. brasilense* also maintains 6 plasmids ranging in size from 0.16 to 1.7 Mb <u>(Orlandini et al. 2014)</u>. The strains designated as biosafety level (BSL) 1 except *K. michiganensis* m5al that harbors gene clusters for the transport of yersiniabactin, a critical virulence factor making it a potential pathogen and a BSL2 organism <u>(Yu et al. 2018)</u>.

Next, we established baseline growth conditions for these non-model microbes for laboratory cultivation. The endophytic bacterial species (*K. michiganensis, A. brasilense,* and *H. seropedicae*) were cultured in standard Luria-Bertani (LB) media. In contrast, root nodule symbionts *R. leguminosarum,* and *A. caulinodans* show more robust colony formation and higher revival rates from cryostorage in tryptone-yeast extract (TY) media (see **Materials and Methods**) suggesting the reduced salt concentration and ionic strength was beneficial for enhanced laboratory growth in several of these species. Growth assays were performed for the endophytic bacteria (*K. michiganensis, A. brasilense,* and *H. seropedicae)* as they readily grew in 96-well plate format. *K. michiganensis* is the fastest growing diazotroph we tested with a doubling time of 26.1 minutes. *H. seropedicae* and *A. brasilense* grew slower with a doubling time of 53 and 87.9 minutes, respectively (**Table 1 & Supplemental Figure 1**).

**Table 1.**
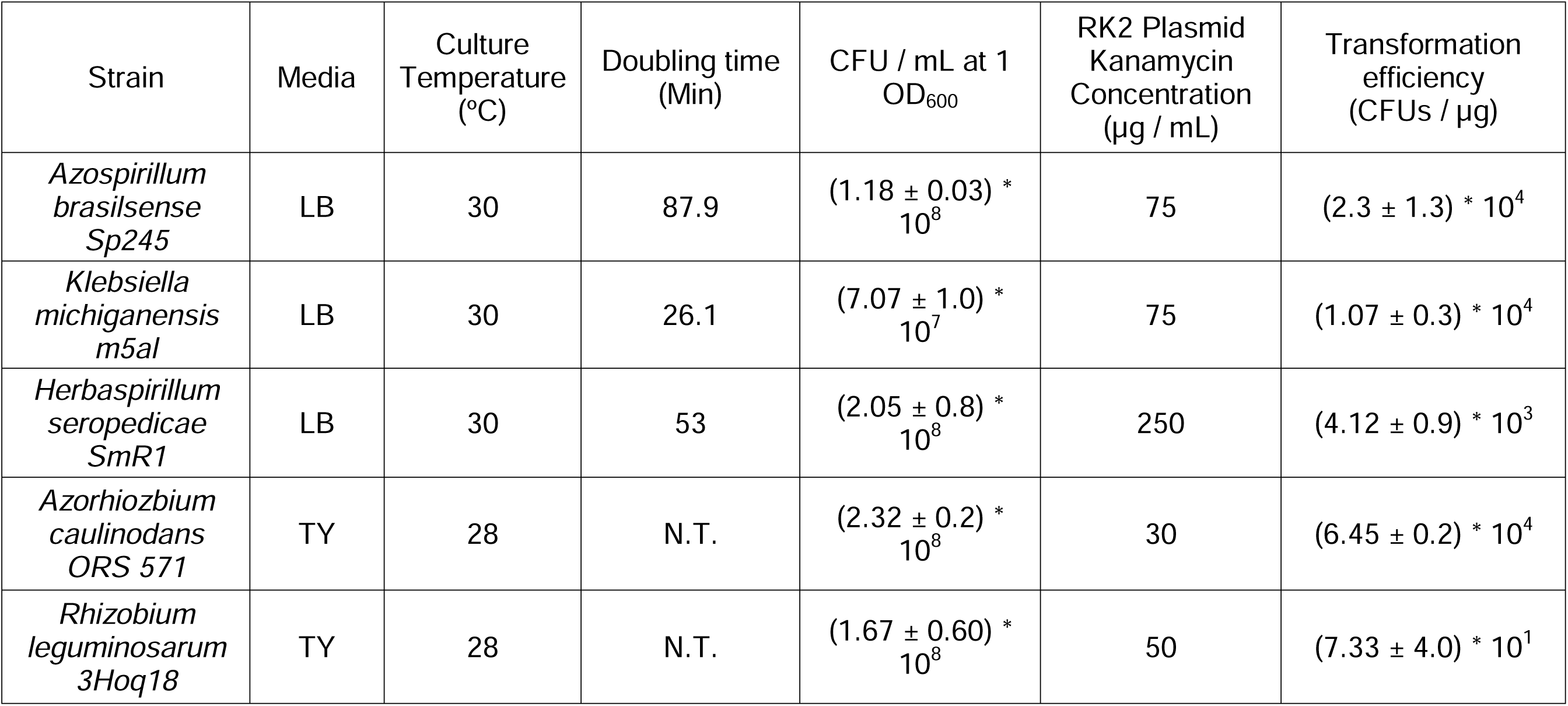
Characterization of selected diazotrophs in this study. From left to right: diazotroph strain, growth media (LB or TY). Standard culture temperature. Doubling time in 96-well format (N.T.: not tested, strains failed to grow in 96-well plate format) (**Supplemental Figure 1**). Colony formation units (CFU) per 1 mL normalized to 1 OD_600_ in log phase (OD_600_ ∼0.5-0.8). Kanamycin selection concentrations for each strain given as µg per mL for plasmids with origin of replication RK2. Transformation efficiencies (CFUs / µg) of each strain for plasmid TE554 harboring RK2 origin of replication (**Supplemental Figure 2**). CFU/mL and transformation efficiency numbers represent three biological replicates and standard deviations are given as error.

Classical mutant analysis studies in diazotrophs rely on time-consuming homologous recombination-based methods for integrating plasmid vectors in species such as *Azotobacter vinelandii* <u>(Dos Santos 2019)</u>. Therefore, we wanted to express genes from stable, exogenously provided plasmid DNA as it enables the rapid characterization of genetic elements and expression of heterologous genes. We first identified the minimum inhibitory concentrations (MIC) for chloramphenicol (**Supplemental Table 2**) and kanamycin (**Table 1**) for each diazotroph with three broad host plasmids BBR1 <u>(Antoine and Locht 1992)</u>, RSK1010 <u>(Scherzinger et al. 1984)</u>, and RK2 <u>(Thomas 1981)</u>. Plasmids with origin of replication RK2 and kanamycin selection marker were stably replicated in five diazotrophic strains, facilitating the characterization of common genetic elements in parallel. Two additional diazotrophs (*Sinorhizobium meliloti 1021* and *Gluconacetobacter diazotrophicus Pal5)* were not readily transformed with any plasmids regardless of electroporation protocol implemented and as a result were not characterized further.

Many gene editing techniques require the introduction of linear DNA fragments or ssDNA (e.g., CRISPR or recombineering) ruling out the possibility of conjugative approaches with helper microbes using *tra-*family conjugation elements, which cannot target these DNA species for introduction into other microbes. Specifically, the ∼90mer ssDNA oligonucleotides designed per targeted edit with recombineering cannot contain the *oriT* sequence required to initiate transconjugation since this additional DNA sequence would introduce a region of non-homology that blocks base-pairing with the targeted locus <u>(Couturier et al. 2023)</u>. Therefore, we constructed small expression plasmids (∼4 kb) each containing an origin of replication and antibiotic selection marker to determine transformation efficiency via electroporation (see **Materials and Methods**). **Supplemental Table 3** summarizes the plasmids used in this study. Plasmids containing the origin of replication BBR1 or RSF1010 were not reliably transformed into *A. brasilense* and *R. leguminosarum* respectively. BBR1 and RSF1010 also yielded high day-to-day variation, and were found to be low efficiency in some species (**Supplemental Table 2,** & additional data not shown). In contrast, plasmids containing the origin of replication RK2 and a kanamycin resistance marker were most widely successful across diazotroph strains. Of the strains that could be successfully transformed, the transformation efficiencies for plasmids containing the origin of replication RK2 were found to be consistent in these four diazotrophs (**Table 1**) exceeding ∼4 x 10^3^ colony formation units (CFU) per µg DNA. Extraction and reintroduction of the same plasmid isolated from *A. brasilense* resulted in an increase of ∼1 order of magnitude of the transformation efficiency, likely due to specific plasmid DNA methylation from replication in *A. brasilense,* preventing degradation as previously described <u>(Mermelstein and Papoutsakis 1993)</u>. While these transformation efficiencies are still several orders of magnitude lower than that of *E. coli*, it is comparable in efficiency to other soil microbes like *Corynebacterium glutamicum* and *Bacillus subtilis* and sufficient for building mutant libraries <u>(Ruan et al. 2015; Xue et al. 1999; Inoue et al. 1990)</u>. Despite the relatively lower efficiencies of the RK2 plasmids in *R. leguminosarum,* this electroporation protocol was reproducible with consistent transformation efficiency across several trials spanning several years and operators. Therefore, we were able to minimize the variety of plasmids used to enable transformation across multiple diazotroph species.

### Genetic Elements for Constitutive Heterologous Protein Expression

Our understanding of gene expression in non-model bacteria remains incomplete; envisioned edits to cellular metabolism often have unpredicted phenotypic behavior <u>(Wintermute and Silver 2010; Banerjee et al. 2024; Rivier et al. 2023)</u>. As a first step in building tunable gene expression platforms, we needed to assess the activity of existing promoters and other genetic elements to determine if they were compatible with diazotrophs, which have divergent sigma factors and transcriptional machinery compared to *E. coli <u>(Zhan et al. 2016)</u>*. To enable reliable dynamic expression, we leveraged the Anderson promoter library (Anderson, 2016). Initially created in *E. coli*, the Anderson promoter library has mutations to conserved regulatory motifs at the –35 and –10 upstream locations in a highly conserved short promoter sequence known to bind transcription factors with the general from 5’-NNNNNNGCTAGCTCAGTCCTAGGNNNNNNGCTAGC-3’. The result is a library of constitutive promoters with discrete transcription rates. We initially tested the consensus promoter from the Anderson collection (J23119) to confirm reliable constitutive expression of an sfGFP fluorescent reporter (**Supplemental Figure 2**). We next constructed a plasmid library with a subset of Anderson promoters to modulate transcription (**Figure 2A**), selecting promoters that exhibited a range of fluorescence expression in *E. coli* as a proxy for transcriptional induction and measured fluorescence of all samples 24 hours post induction. The representative library resulted in changes of expression of up to 100-fold in *K. michiganensis* and consistent expression changes of ∼10-fold in the remaining diazotroph species compared to the consensus sequence (J23119) (**Figure 2B-F**). At best, the consensus promoter J23119 had at least 5-fold or higher expression than the J23103 promoter as observed in *E. coli*, but overall trends were harder to discern. When Anderson promoter variants were compared across diazotrophs, the rank order of fluorescence expression was not conserved. Moreover, their ranked expression level was also different across each microbe, highlighting the necessity to test each expression element (**Supplemental Figure 4**).

**Figure 2.**
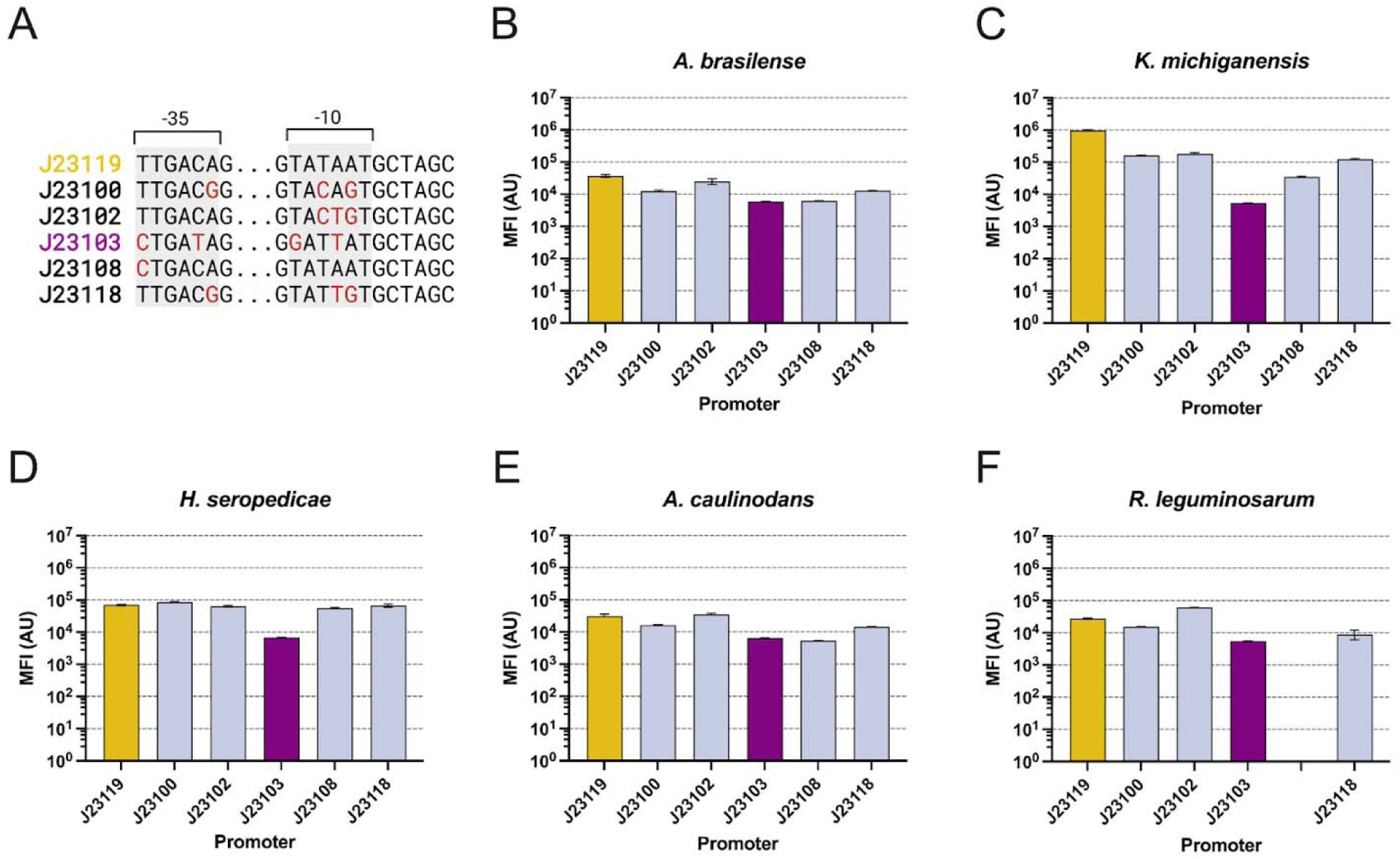
Expression of sfGFP using Anderson Promoters. Schematic of the Anderson promoter library (A) highlighting nucleotide differences (red) from wildtype promoter J23119 (yellow) and the lowest expression promoter as assessed in *E. coli* J23103 (purple). Mean fluorescence intensity (MFI) in arbitrary units (AU) across selected diazotrophs in logarithmic scale (B-F). Yellow and purple bars represent the most highly and lowly expressed promoters in *E. coli*, respectively. Promoters are organized from highest fluorescence to lowest for each diazotroph (left to right). Error bars represent standard deviation over three biological replicates. Full promoter sequences can be found in Supplemental table S2.

After modulating expression by varying promoter sequences, we next tested a library of ribosomal binding sites (RBS) to modulate translation. We built an RBS library using the De Novo DNA platform consisting of eleven variable RBSs predicted to yield a range of translation rates in *A. brasilense* <u>(Reis and Salis 2020; Farasat et al. 2014; Ng et al. 2015; Espah Borujeni et al. 2014, 2017)</u>. The resulting degenerate DNA RBS library genetic sequence (5’ GTGYASAGGCAARAARGASGTHTTTA 3’) was varied at 6 nucleotide locations and predicted to have a range of expression of 1000-fold in *A. brasilense*. Full RBS sequences are summarized in **Supplemental Table 4**. The RBSs were then inserted into plasmids containing an mScarlet reporter along with the consensus Anderson promoter J23119 (**Figure 3A**). The resulting *A. brasilense* optimized RBS-variant plasmids were examined for mScarlet fluorescence in all five microbes. We observed that the relative mScarlet fluorescence varied across all bacterial species, further enabling dynamic expression (**Figure 3B-F**). The observed change in *A. brasilense* was ∼10-fold and followed the general trend as computationally predicted. Larger ranges of expression were noted for several species including a ∼100-fold range in *H. seropedicae* and *A. caulinodans*. However, the other four diazotrophs followed no correlation with the predicted values, underscoring the difference between bacterial strains (**Figure 3C-F)**. The predicted strongest (**Figure 3C-F, light blue)** and lowest (**Figure 3B-F, pink)** are not consistent across strains and do not constitute the experimentally determined strongest and weakest RBSs as predicted in *A. brasilense* **(Figure 3D).** Moreover, the normalized relative fluorescence was not conserved across bacterial strains similar to the promoter sequences, again highlighting the need to test each individual component and tailor RBS libraries for each microbe (**Figure 3G**). Several RBS plasmid variants failed transformation in several species, but the reason for this limitation is unknown. A second, smaller RBS library was built using the same predictive platform in *A. brasilense* to control expression of sfGFP offering a library of distinct RBS sequences (**Supplemental Figure 5A**). The resulting library shows less dynamic range (∼10-fold) than the previous library but still could offer additional expression control given the variability of downstream applications or target genes (**Supplemental Figure 5B-F**). Given the small library size relative to previous studies in *E. coli*, testing a larger RBS sample space (e.g., ∼500 RBS variants) could identify a wider expression range in diazotrophs <u>(Espah Borujeni and Salis 2016)</u>.

**Figure 3.**
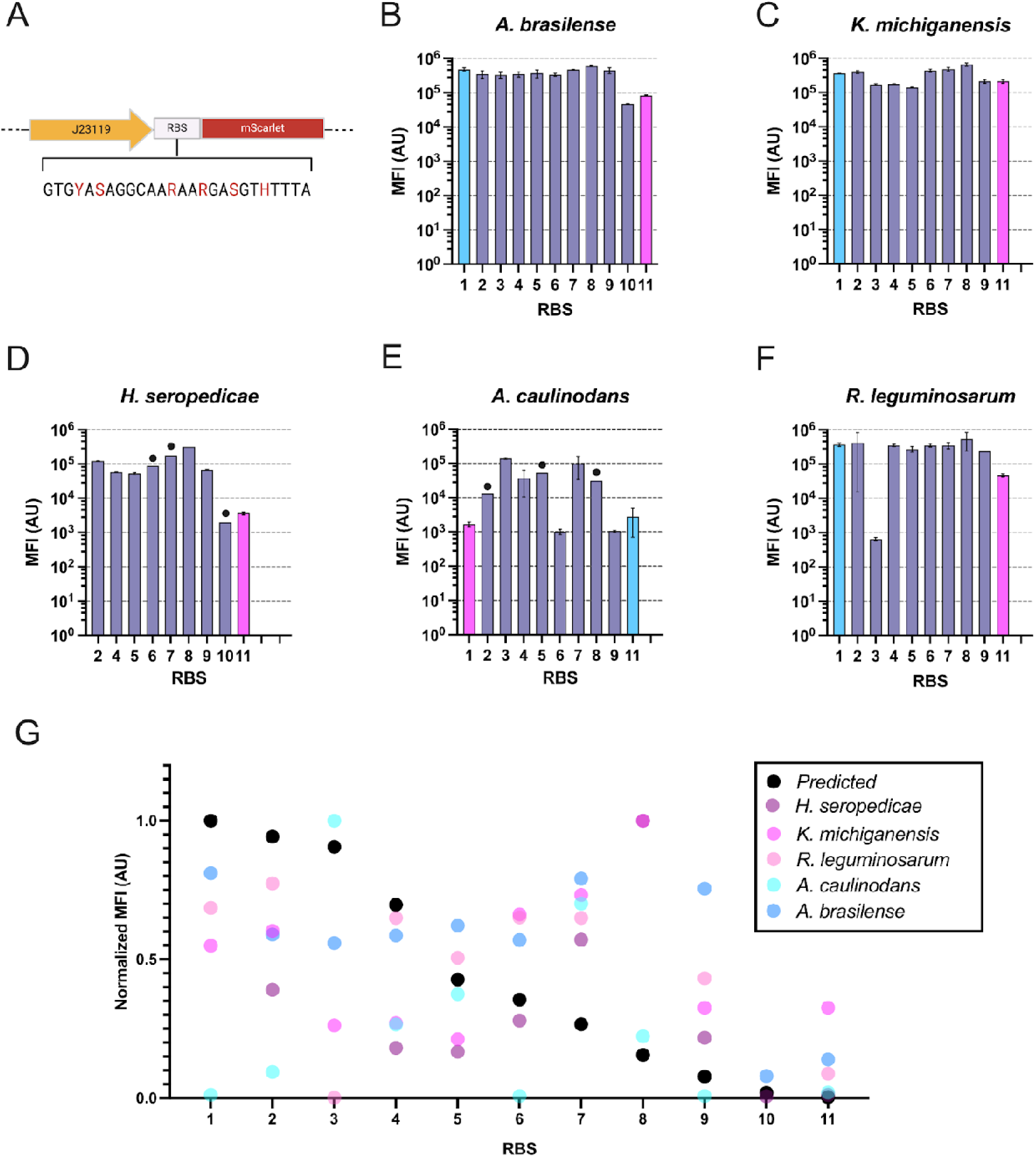
Behavior of *A. brasilense*-optimized Ribosomal binding sites on gene expression acros divergent diazotrophs. Representation of RBS plasmid library harboring mScarlet fluorescence protein (A) controlled by J23119 promoter (yellow) and variable RBSs by altering nucleotide bases highlighted in red. Mean fluorescence intensity (MFI) in arbitrary units (AU) of mScarlet in logarithmic scale for selected diazotrophs (B-F). RBSs are organized by decreasing fluorescence in each diazotroph. Light blu and pink bars represent the RBS to have the highest and lowest predicted activity, respectively. Error bars represent standard deviation over three biological replicates. Dots denote one biological replicate. Full RBS sequences are available in **Supplemental Table 2**. Highest fluorescence normalized to a value of one reveals divergence of RBSs in different species (G).

By testing a library of common genetic expression elements, we were able to test five strains in parallel. However, we found varying efficacies to modulate expression at the transcriptional and translational level. For example, the promoters tested for *K. michiganensis* result in a wider range of expression (∼100-fold) compared to the RBSs (<10-fold), but the opposite is true for *A. caulinodans* and *R. leguminosarum* (**Figure 3C, E, F**). Nonetheless, our approach facilitated rapid parallel characterization of multiple strains and resulted in detectable fluorescence in all diazotrophs. The ability to modulate expression is crucial for the establishment of synthetic genetic networks, especially in diazotroph-plant or diazotroph-microbe interactions where signaling molecule interactions can vary across orders of magnitude <u>(Cesco et al. 2010)</u>. Considerable effort has gone into further understanding how to more effectively create sustained microbe-diazotroph associations that result in higher amounts of ammonia for the plant. The ability to express exogenous DNA expands our capacity to investigate specific genes involved in these complex associations. Moreover, genetic circuits between plants and soil bacteria in the rhizosphere have been targets for monitoring and controlling plant physiology. The tools demonstrated here offer potential to leverage diazotrophic bacteria to modulate diazotroph strain behavior and affect their environment by the expression of heterologous genes, perhaps by changing the composition of the plant microbiome with new secreted metabolites or proteins.

### Inducible Gene Expression Systems in Diazotrophs

In addition to the constitutive expression system generated above, conditional expression of heterologous genes is a desirable tool for the construction of synthetic circuits especially when potentially growth-inhibitory genes are introduced. Therefore, we tested several small molecule inducible systems previously optimized in the “Marionette” *E. coli* strain <u>(Meyer et al. 2019)</u>. We specifically chose naringenin, arabinose, salicylic acid, and vanillic acid, which are inducer molecules that are also known to exist in root exudates <u>(Vives-Peris et al. 2020)</u>. The promoter sequences for these inducible systems were subcloned into existing plasmid designs to drive expression of sfGFP. After construction, these plasmids were transformed into the selected diazotrophs and, as in *E. coli*, 0.05 μM to 2000 μM inducer was added <u>(Meyer et al. 2019)</u>. We found the most robust induction with salicylic acid, yielding over a 10,000x increase in expression compared to the uninduced control in *H. seropedicae* (**Figure 4A**), *K. michiganensis* (**Figure 4B**), and *R. leguminosarum* (**Figure 4C**). In addition, arabinose induced expression in *K. michiganensis* and vanillic acid had limited effect in *R. leguminosarum* and only showed modest fold increases at millimolar concentrations. None of these promoter systems were portable into *A. brasilense* and *A. caulinodans* and would require additional optimization to determine the rate limiting step blocking pathway induction since we the fluorescent output was previously validated with a constitutive promoter system (**Supplemental Figure S6,** refer to **Figures 3 & 4**). It is possible that root exudate compounds may necessitate additional consideration as they may already exist in native diazotroph metabolism as suggested by the degradation of naringenin in *H. seropedicae* <u>(Marin et al. 2013)</u>. Regardless, by establishing inducible systems in diazotrophs, we enabled a prototypical synthetic gene circuit for exploring plant-microbe interactions responsive to the temporal kinetics found in plant exudate. Regardless, by establishing inducible systems in diazotrophs, we enabled a prototypical synthetic gene circuit for exploring plant-microbe interactions responsive to the temporal kinetics found in plant exudate.

**Figure 4.**
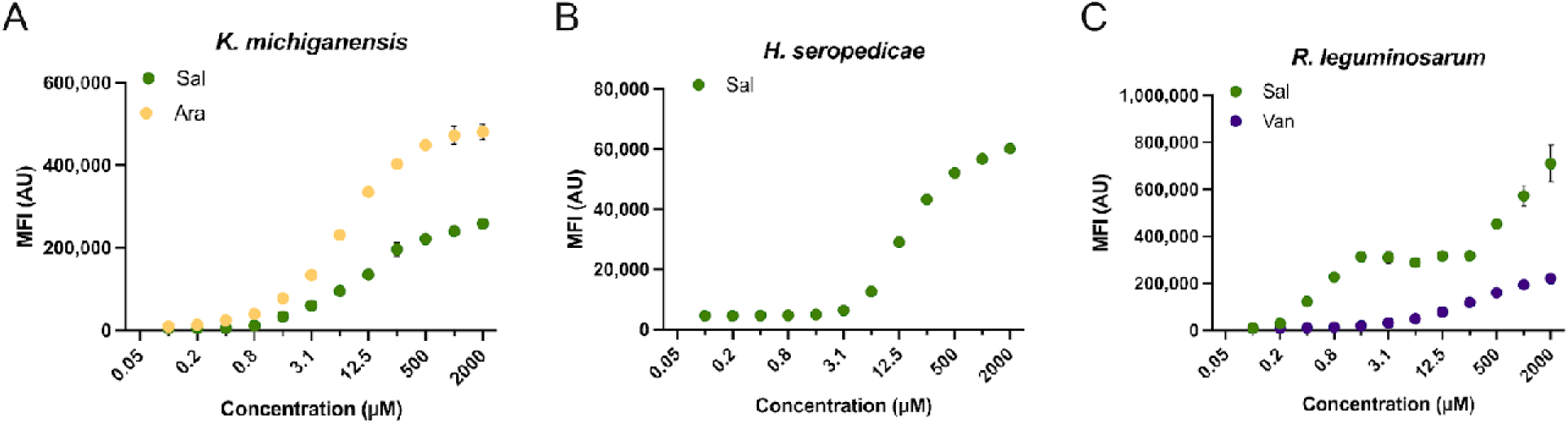
Inducible promoter systems in diazotrophs. Mean fluorescence intensity (MFI) in arbitrary units (AU) of inducible promoter systems in *H. seropedicae* (A), *K. michiganensis* (B), and *R. leguminosarum* (C) plotted in linear scale. Small molecule inducers arabinose (Ara, yellow), salicylic acid (Sal, green), and vanillic acid (Van, purple) were varied in concentration from 0.05 μm to 2000 μm. Error bars represent standard deviation over two biological replicates. Non-functional inducer systems are summarized in **Supplemental Figure 7**

The expression systems in diazotrophs described here may be further leveraged to improve diazotroph ammonia release or plant associations. Genetic components of nitrogenase related genes could be expressed in a tailored fashion, potentially enabling scenarios where an increase in exogenous ammonia can be observed as has been previously reported in other diazotrophs <u>(Mus et al. 2022)</u>. Moreover, plant associations could be engineered to increase diazotroph ability to establish in plant roots. Increased plant interaction could be fostered through the over-expression of genes known to be involved with endophytic relationships <u>(Cole et al. 2017)</u>. Likewise, gene expression of biocontrol traits could offer methods for engineered strains to outcompete other bacterial species, including pathogens, and result in increased plant associations. Regardless of the downstream approach, coupling these expression-based tools with other methods such as adaptive evolution may be necessary given the complex nature of these biological systems <u>(Nordgaard et al. 2022)</u>.

### Demonstration of Genome Editing in K. *michiganensis*

Having developed a broad set of plasmids compatible with electroporation methods, we proceeded to test targeted genome editing using a plasmid-borne recombineering system in conjunction with a chemically synthesized oligonucleotide <u>(Datta et al. 2008)</u>. By overexpressing components of the mismatch repair system containing a dominant negative MutL-E36K mutation, the provided oligo initiates lagging strand DNA replication, incorporating the desired genomic mutation <u>(Fernández-Cabezón et al. 2021)</u>. To apply this method in a diazotroph, we used an oligo targeting the ribosomal 12S gene (*rpsL*) that confers streptomycin resistance by introducing a Lys to Arg mutation at amino acid residue 43 with three mutated nucleotides <u>(Wannier et al. 2020)</u>. We picked *K. michiganensis* for this demonstration because it has the fastest doubling time and formed colonies after plasmid transformation within 2 days of plating. When this oligonucleotide is electroporated into cells expressing both *recT* and *mutL-E36K* from *Pseudomonas aeruginosa,* we expect to see an increase in streptomycin resistant colonies compared to a control reaction. Consistent with this model, electroporation of *K. michiganensis* with this oligonucleotide yielded ∼10 streptomycin resistant colonies with no streptomycin resistant clones without addition of the oligonucleotide. Several randomly chosen streptomycin resistant clones were sequenced at the *rpsL* locus and all were confirmed to harbor the expected K43R point mutation through Sanger sequencing where the identical mismatched nucleotides chemically synthesized in the oligonucleotide were incorporated into the *K. michiganensis* genome (**Supplemental Figure 7**). These results demonstrate that other genomic edits at other loci can be generated with other selection markers (e.g., auxotrophies or antibiotics) and potentially extended with the development of targeted endonuclease methods like CRISPR for scarless modifications.

## Conclusion

Here, we have outlined the growth and cultivation parameters for five diazotrophs in laboratory conditions and identified a suite of genetic elements. Specifically, we attempted to onboard seven phylogenetically distinct diazotrophs finding five suitable for exogenous DNA expression. We then tested eleven constitutive promoters and a total of twenty RBSs between two libraries. The tested genetic expression elements enable translation and transcription attenuation of up to ∼100-fold and ∼1000-fold, respectively. Additionally, four root exudate compounds were tested for the efficacy in induction resulting in five systems of gene induction across three diazotrophs. A larger set of genetic tools will enable other researchers in the field to manipulate more components of rhizobial community members with the potential to improve bioenergy crop yields while reducing dependence on energetically expensive and environmentally detrimental fertilizers.

## Materials and Methods

### Bacterial strains, growth conditions, and cell counts

Bacterial strains *Herbaspirilium seropedicae* SmR1, *Klebsiella michiganensis* M5al, *Azospirillum brasilense* Sp245 were generously donated by Adam Deutschbauer. *Azorhizobium caulinodans* ORS571, and *Rhizobium leguminosarum* 3D1k2 were obtained from The American Type Culture Collection (ATCC) and are described in the strain table (**Table 1**). All microbes used in this study were verified with 16s rRNA sequencing using primers 27F (forward: 5′-AGAGTTTGATCMTGGCTCAG-3′) and 1492R (reverse: 5′-TACGGYTACCTTGTTACGACTT-3′) <u>(Weisburg et al. 1991)</u>. All strains were stored in 10% glycerol at –80 °C. Plasmid sequences are available at public-registry.jbei.org after generating a free user account and plasmids are available upon request to the corresponding author.

*A. Brasilense*, *K. michiganensis*, and *H. seropedicae* were cultured in autoclaved LB media (Tryptone, 10.0 g/L; Yeast extract, 5.0 g/L; NaCl, 10.0 g/L) at a temperature of 30 °C. *A. caulinodans* was cultured in autoclaved TY media (Tryptone 5.0 g/L, Yeast extract 3.0 g/L, CaCl_2_ x 2 H_2_O 0.9 g/L) at 28 °C. Tryptone, yeast extract, NaCl, and CaCl_2_ were purchased from BD Biosciences (Milpitas, CA).

Kinetic growth curves were generated by measuring optical density (OD) at a wavelength of 600 nm and pathlength of 1 cm every 10 minutes using a Molecular Devices Filtermax F5 plate reader (Molecular Devices LLC, San Jose, CA). Microtiter plates were incubated at 30 °C for the duration of the time course at “high” shake speed setting and linear shaking mode in a 96 well microtiter plate sealed with a Breathe-Easylll transparent membrane (Sigma-Aldrich). Three biological replicates were processed by growthcurver R package and doubling times were found for all strains <u>(Sprouffske and Wagner 2016)</u>.

Colony formation unit of bacterial strains were measured by growing each strain in their respective media (**Table 1**) for ∼16-24 hours to logarithmic growth except for the fast-growing strain *K. michiganensis* that was grown for 16 hours then back diluted to an OD_600_ of 0.1 and grown for ∼2 hours to logarithmic phase. Cells were then diluted in phosphate buffered saline (PBS, Thermofisher scientific, Waltham, MA) to a range of dilutions (10,000x – 2,000,000x) and plated onto LB agar or TY agar plates and incubated at 30°C for 2-5 days until colonies could be visualized. Dilution ranges were selected for calculating initial OD to CFU values where roughly 200 CFUs per plate <u>(Tomasiewicz et al. 1980)</u>. Three biological replicates of CFUs were counted and back calculated to the original concentration and normalized to an OD_600_ of 1.

### Plasmid Construction

All DNA constructs and oligos were designed using either Snapgene (BioMatters Ltd) or Primer3 <u>(Koressaar and Remm 2007)</u> and plasmids were assembled using isothermal HiFi assembly (New England Biolabs (NEB), Ipswitch, MA) following manufacturer’s guidelines. Ribosomal binding site libraries were predicted using the De Novo DNA platform with either mScarlet or GFP as the protein coding sequence and *A. brasilense* as the host organism. DNA amplification was conducted following manufacturer’s guidelines. Whole plasmid sequencing (Primordium Labs, Monrovia, CA) was used to verify constructs as necessary. Standard *E. coli* bacterial transformation with Invitrogen XL-1 Blue competent cells (ThermoFisher, Carlsbad, CA) was used to isolate new DNA constructs. Plasmid DNA was isolated using alkaline lysis and silica columns using Qiagen miniprep kits (Qiagen Inc, Germantown MD) following the manufacturer’s guidelines.

### Bacterial transformations

Bacterial strains were grown in 5 mL of the appropriate media (**Supplemental Table 1**) for 24 hours at 30 °C to reach saturation. Cultures were diluted to an OD_600_ of 0.1 in 25 mL of liquid media in a 250 mL flask and incubated until they reached an OD_600_ of ∼0.5 (approximately ∼2-6 hours). *R. leguminosarum* was notably slow growing and was diluted to an initial OD_600_ of 0.05 and incubated overnight to reach a comparable OD_600_ in logarithmic phase by the next morning. Cells were harvested by centrifugation at 5000 x*g* for 10 minutes and washed twice with ice-cold 10% (v/v) glycerol. The pellets were resuspended in 250 μL of 10% glycerol and 50 μL aliquots were prepared and stored at –80 °C. Cell aliquots were thawed on ice for 5 minutes before adding 100 ng of plasmid DNA and electroporated on Bio-rad Micropulser (Bio-Rad, Hercules, CA) using EC2 protocol (0.2 cm cuvette, 2.5 kV). This protocol was modified slightly for *R. leguminosarum* by resuspending 4 mL of culture media in 100 μL of 10% glycerol to increase the number of cells used per electroporation. Resulting transformants were then grown in liquid media for 5 hours and plated onto respective selection plates with appropriate antibiotic concentrations (**Supplemental Table 1**).

Transformation efficiency numbers were calculated by harvesting cell cultures (grown as described above) to logarithmic phase. Cells were diluted to an OD_600_ of 0.1 in PBS pH 7.4, then serial diluted to low concentrations (OD_600_ of 1.0 x 10^-6^, 1.0 x 10^-5^) and plated on respective agar plates. Colony formation units (CFU) were counted and normalized to account for dilutions to determine equivalent CFUs for 1 unit of OD.

### ssDNA Recombineering in *K. michiganensis*

ssDNA recombineering was performed in *K. michiganensis* with slight modifications from our existing recombineering protocol used in *Pseudomonas putida <u>(Czajka et al. 2022)</u>.* The strain was transformed with a plasmid containing a 3-methyl benzoate (3MB; m-Toluic Acid; Sigma Aldrich Catalog No. T36609) inducible recT recombinase and *mutL^E36K^* (pTE583) was grown overnight, back diluted to a starting OD_600_ of 0.1, and allowed to grow for an additional 3 hours. The recombineering pathway genes were induced with 1 mM 3MB for 4 hours before being electroporated as described with oligonucleotide containing a K43R triple point mutation (5’ GCAGAAACGTGGCGTATGTAC TCGTGTATATACCACCACTCCTCGTAAACCTAACTCCGCACTGCGTAAAGTTTGCCGTGTG CGTCTGAC 3’) (Integrated DNA Technologies, Redwood City, CA). The oligo was ordered desalted and without further purification or base modifications. Resulting transformants were screened on solid LB agar media containing 100 µg/mL streptomycin (Sigma Aldrich). Transformants were confirmed to incorporate the identical three nucleotide polymorphism by Sanger sequencing (**Supplemental Figure 7**). No spontaneous streptomycin clones were recovered on the control transformation plates without electroporation with oligo.

### Quantifying fluorescent expression

Flow cytometric analysis was conducted using an Accuri C6 flow cytometer equipped with an autosampler (BD Biosciences). Cells were cultivated in 1 mL media in 24 well deep well plates at 30 °C and 250 rpm for 24 hours prior to being diluted to OD_600_ 0.1 in 500 μl of the respective medium in a 96 well microtiter plate for fluorescence measurements. Measurements were conducted in biological triplicate. A total of 30,000 events were recorded at the “low” flow rate setting with a core size of 22 μm. GFP was excited at 478 nm at 70 mW and emission detected at 530 nm.

Vanillic acid, arabinose, and salicylic acid were dissolved in sterile deionized water. Naringenin was dissolved in dimethyl-sulfoxide (DMSO). Expression was induced via small molecules by incubating for 24 hours and induction of mScarlet was excited at 552 nm with default signal intensity and emission detected at 610 nm. For all fluorescent measurements, data acquisition was performed as described in the Accuri C6 Sampler User’s Guide. The acquired data were analyzed in FlowJO (TreeStar Inc, San Carlos, CA).

## Author Contributions

Molecular biology and strain engineering: SK, AR, SM, TE. Data interpretation and analysis: SK, AR, RK, TE. Supervision: TE. Drafted the manuscript: AR, TE. Acquisition of Funds: TE. All authors have read, provided feedback, and approved the manuscript for publication.

## Acknowledgements

We thank Adam Deutschbauer for the kind gift of several nitrogen fixing bacteria used in this study, and members of the A. Mukhopadhyay and A. Eudes groups for their constructive feedback during this project. We thank David Carruthers and Aindrila Mukhopadhyay (LBNL) for feedback and suggestions on the draft manuscript. We acknowledge technical assistance from Jess Sustarich and William Gaillard (Sandia National Lab) in developing transformation and cultivation protocols, and Nathan Hillson for guidance and leadership as part of the DOE Agile BioFoundry project. Graphical icons used in Figures in this study were generated in BioRender (Science Suite Inc) and graphs were generated in GraphPad Prism 10 (Insight Partners LLC).

