## Supplemental Figures for "Development of Modular Expression and Genome Editing Across Phylogenetically Distinct Diazotrophs"

**Supplemental information**

| **Strain** | **Origin** | **Climate** | **Reference** | **ATCC** | **DSMZ** | **NCBI Genome accession number** | **GC content (%)** | **Genome size (Mb)** | **Chromosomes** |
| --- | --- | --- | --- | --- | --- | --- | --- | --- | --- |
| *Azospirillum brasilense* Sp245 | Paraná | humid subtropical | [(Baldani et al., 1987)](https://sciwheel.com/work/citation?ids=16357304&pre=&suf=&sa=0) | BR-11005 | N.A. | GCF_003119195.2 | 68.5 | 7.7 | 1 (6 plasmids) |
| *Klebsiella michiganensis* m5al | Unknown | Unknown | [(Pengra and Wilson, 1958)](https://sciwheel.com/work/citation?ids=16357323&pre=&suf=&sa=0) | BAA-1236 | 7342 | GCA_000308735.2 | 56 | 5.8 | 1 |
| *Herbaspirillum seropedicae* SmR1 | Rio de janiero | tropical savanna | [(Baldani et al., 1986)](https://sciwheel.com/work/citation?ids=16102296&pre=&suf=&sa=0) | 35893 | 6445 | GCF_000143225.1 | 63.5 | 5.5 | 1 |
| *Azorhiozbium caulinodans* ORS 571 | Senegal | Tropical | [(Dreyfus et al., 1988)](https://sciwheel.com/work/citation?ids=16357330&pre=&suf=&sa=0) | 43989 | 5975 | GCF_000010525.1 | 67.5 | 5.4 | 1 |
| *Rhizobium leguminosarum* 3Hoq18 | America (non-descript) | Unknown | [(Bergersen, 1958)](https://sciwheel.com/work/citation?ids=16357333&pre=&suf=&sa=0) | 14479 | 6040 | GCF_033842645.1 | 60.5 | 5.1 | 1 |

**Supplemental Table 1.** Strains used in this study. Background information including origin, references to climate, repository accession numbers and genome details.

| **Strain** | **RSF1010** | **BBR1** | **RK2** | **Kanamycin Concentration (μg / mL)** | **Chloramphenicol Concentration (μg / mL)** |
| --- | --- | --- | --- | --- | --- |
| *Azospirillum brasilense Sp245* | ✓ | X | ✓ | 75 | 100 |
| *Klebsiella michiganensis m5al* | ✓ | ✓ | ✓ | 75 | 100 |
| *Herbaspirillum seropedicae SmR1* | ✓ | ✓ | ✓ | 250 | 200 |
| *Azorhiozbium caulinodans ORS 571* | ✓ | ✓ | ✓ | 30 | 100 |
| *Rhizobium leguminosarum 3Hoq18* | X | ✓ | ✓ | 50 | 100 |
| *Gluconacetobacter diazotrophicus Pal5* | X | X | X | N/A | N/A |
| *Sinorhizobium meliloti 1021* | X | X | X | N/A | N/A |

**Supplemental Table 2.** List of origins of replications attempted to use for transformation by electroporation across bacterial species in this study. Check mark denotes successful transformation, X denotes no colonies obtained.

| Plasmid | Origin of Replication | Marker | Description | Joint Bioenergy Institute repository number |
| --- | --- | --- | --- | --- |
| pTE554 | RK2 | Kanamycin | Empty - |  |
| pTE577 | RK2 | Gentamicin | mCherry |  |
| pTE583 | RK2 | Gentamicin | Pap_RecT Pa_MutL-E36K | JBEI-258211 |
| pTE697 | RK2 | Chloramphenicol | empty |  |
| pTE698 | RK2 | Chloramphenicol | J23119-mCherry |  |
| pTE717 | RK2 | Chloramphenicol | J23119-mCherry |  |
| pTE739 | RK2 | Kanamycin | J23119-mCherry |  |
| pTE740 | RK2 | Kanamycin | J23119-RBS-mCherry |  |
| pTE741 | RK2 | Kanamycin | RBS-mCherry |  |
| pTE779 | BBR1 | Kanamycin | N/A |  |
| pTE780 | RSF1010 | Kanamycin | N/A |  |
| pTE781 | RK2 | Kanamycin | J23119-sfGFP |  |
| pTE782 | RK2 | Kanamycin | J23119-RBS-sfGFP |  |
| pTE783 | RK2 | Kanamycin | J23119-spacers-RBS-mCherry |  |
| pTE794 | RK2 | Kanamycin | J23119-sfGFP-RBS-012 |  |
| pTE795 | RK2 | Kanamycin | J23119-sfGFP-RBS-013 |  |
| pTE796 | RK2 | Kanamycin | J23119-sfGFP-RBS-014 |  |
| pTE797 | RK2 | Kanamycin | J23119-sfGFP-RBS-015 |  |
| pTE798 | RK2 | Kanamycin | J23119-sfGFP-RBS-016 |  |
| pTE799 | RK2 | Kanamycin | J23119-sfGFP-RBS-017 |  |
| pTE800 | RK2 | Kanamycin | J23119-sfGFP-RBS-018 |  |
| pTE801 | RK2 | Kanamycin | J23119-sfGFP-RBS-019 |  |
| pTE802 | RK2 | Kanamycin | J23119-sfGFP-RBS-020 |  |
| pTE803 | RK2 | Kanamycin | J23119-sfGFP-RBS-021 |  |
| pTE804 | RK2 | Kanamycin | J23119-sfGFP-RBS-022 |  |
| pTE805 | RK2 | Kanamycin | J23119-sfGFP-RBS-023 |  |
| pTE826 | RK2 | Kanamycin | J23119-mScarlet |  |
| pTE834 | RK2 | Kanamycin | J23119-mScarlet-RBS-01 | JBEI-258213 |
| pTE835 | RK2 | Kanamycin | J23119-mScarlet-RBS-02 |  |
| pTE836 | RK2 | Kanamycin | J23119-mScarlet-RBS-03 |  |
| pTE837 | RK2 | Kanamycin | J23119-mScarlet-RBS-04 |  |
| pTE838 | RK2 | Kanamycin | J23119-mScarlet-RBS-05 |  |
| pTE839 | RK2 | Kanamycin | J23119-mScarlet-RBS-06 |  |
| pTE840 | RK2 | Kanamycin | J23119-mScarlet-RBS-07 |  |
| pTE841 | RK2 | Kanamycin | J23119-mScarlet-RBS-08 |  |
| pTE842 | RK2 | Kanamycin | J23119-mScarlet-RBS-09 |  |
| pTE843 | RK2 | Kanamycin | J23119-mScarlet-RBS-10 |  |
| pTE844 | RK2 | Kanamycin | J23119-mScarlet-RBS-11 |  |
| pTE852 | RK2 | Kanamycin | J23100-sfGFP-RBS-012 |  |
| pTE853 | RK2 | Kanamycin | J23102-sfGFP-RBS-012 |  |
| pTE854 | RK2 | Kanamycin | J23103-sfGFP-RBS-012 |  |
| pTE856 | RK2 | Kanamycin | J23108-sfGFP-RBS-012 |  |
| pTE857 | RK2 | Kanamycin | J23112-sfGFP-RBS-012 |  |
| pTE858 | RK2 | Kanamycin | J23118-sfGFP-RBS-012 |  |
| pTE861 | RK2 | Kanamycin | pTtg-GFP-RBS-012 |  |
| pTE862 | RK2 | Kanamycin | pBAD-GFP-RBS-012 |  |
| pTE863 | RK2 | Kanamycin | pSalTTC-GFP-RBS-012 |  |
| pTE864 | RK2 | Kanamycin | pVanCC-GFP-RBS-012 |  |

**Supplemental Table 3.** List of plasmids used or generated in this study.

| **Oligos** | **Sequence (5’-3’)** | **Description & JBEI Repository Number** |
| --- | --- | --- |
| TEAM-3027 | GCAGAAACGTGGCGTATGTACTCGTGTATATACCACCACTCCTCGTAAACCTAACTCCGCACTGCGTAAAGTTTGCCGTGTGCGTCTGAC | recombineering oligo for K43R substitution conferring spectromycin resistance in *Klebsiella michiganensis m5aI* |
| TEAM-3028 | GGAGCTATTTAATGGCAACAATTAACCAGC | colony PCR primer for *Klebsiella michiganensis m5aI* rpsL recombineering |
| TEAM-3029 | GAGCGAGCTTGCTTACGGTC | colony PCR primer for *Klebsiella michiganensis m5aI* rpsL recombineering |
| RBS-01 | GTGCAGAGGCAAAAAGGAGGTTTTTA | mScarlet, Figure 4 |
| RBS-02 | GTGCAGAGGCAAAAAGGAGGTATTTA | mScarlet, Figure 4 |
| RBS-03 | GTGCACAGGCAAGAAGGAGGTTTTTA | mScarlet, Figure 4 |
| RBS-04 | GTGCACAGGCAAGAAGGAGGTATTTA | mScarlet, Figure 4 |
| RBS-05 | GTGCAGAGGCAAAAAGGAGGTCTTTA | mScarlet, Figure 4 |
| RBS-06 | GTGTAGAGGCAAGAAGGAGGTTTTTA | mScarlet, Figure 4 |
| RBS-07 | GTGTACAGGCAAGAAGGAGGTTTTTA | mScarlet, Figure 4 |
| RBS-08 | GTGTACAGGCAAAAAGGAGGTATTTA | mScarlet, Figure 4 |
| RBS-09 | GTGCACAGGCAAGAAAGAGGTATTTA | mScarlet, Figure 4 |
| RBS-10 | GTGTACAGGCAAAAAGGACGTATTTA | mScarlet, Figure 4 |
| RBS-11 | GTGCAGAGGCAAGAAAGACGTCTTTA | mScarlet, Figure 4 |
| RBS-12 | TTTGTTATGAAAGAAAGGAGGTAAGGG | sfGFP, Supplemental Figure 5 |
| RBS-13 | TTTGTTATGAAAGAAAGGAGGTAATGG | sfGFP, Supplemental Figure 5 |
| RBS-14 | TTTGTTATGAAAGTAAGGAGGTAAGGG | sfGFP, Supplemental Figure 5 |
| RBS-15 | TTTGTTATGAAAGAAAGGGGGTAAGGG | sfGFP, Supplemental Figure 5 |
| RBS-16 | TTTGTTATGAAAGTAAGGAGGTAATGC | sfGFP, Supplemental Figure 5 |
| RBS-17 | TTTGTTATGAAAGAAAGGGGGTAATGC | sfGFP, Supplemental Figure 5 |
| RBS-18 | TTTGTTATGAAAGTAAGGGGGTCATGG | sfGFP, Supplemental Figure 5 |
| RBS-19 | TTTGTTATGAAAGAAAGGAGGCAATGC | sfGFP, Supplemental Figure 5 |
| RBS-20 | TTTGTTATGAAAGAAATGAGGTCATGG | sfGFP, Supplemental Figure 5 |

**Supplemental Table 4.** List of oligos and ribosomal binding sites used in this study.

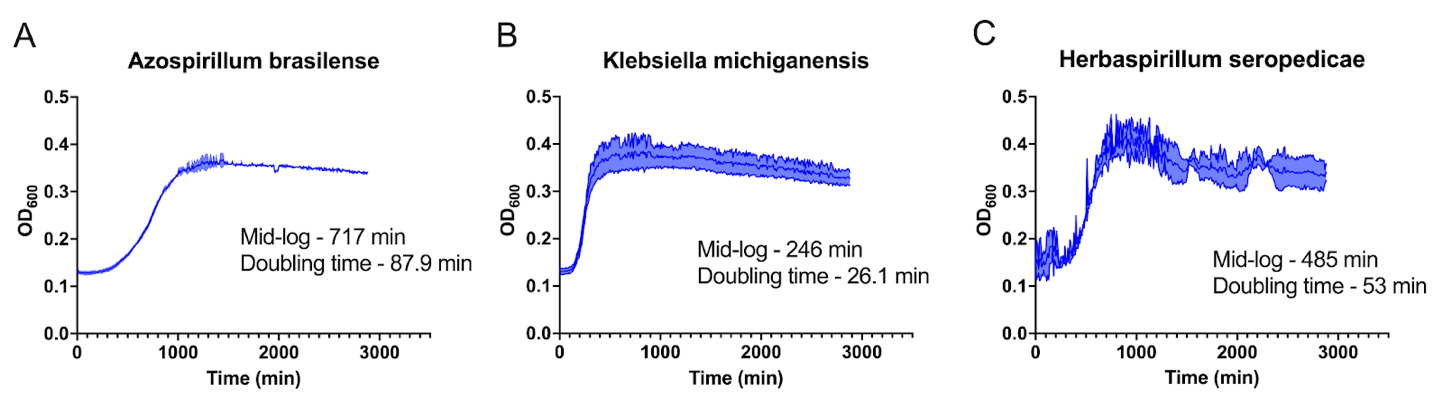

**Supplemental Figure 1. Growth Curves of select Diazotrophs**. 96-well growth curves for the endophytic diazotroph strains.

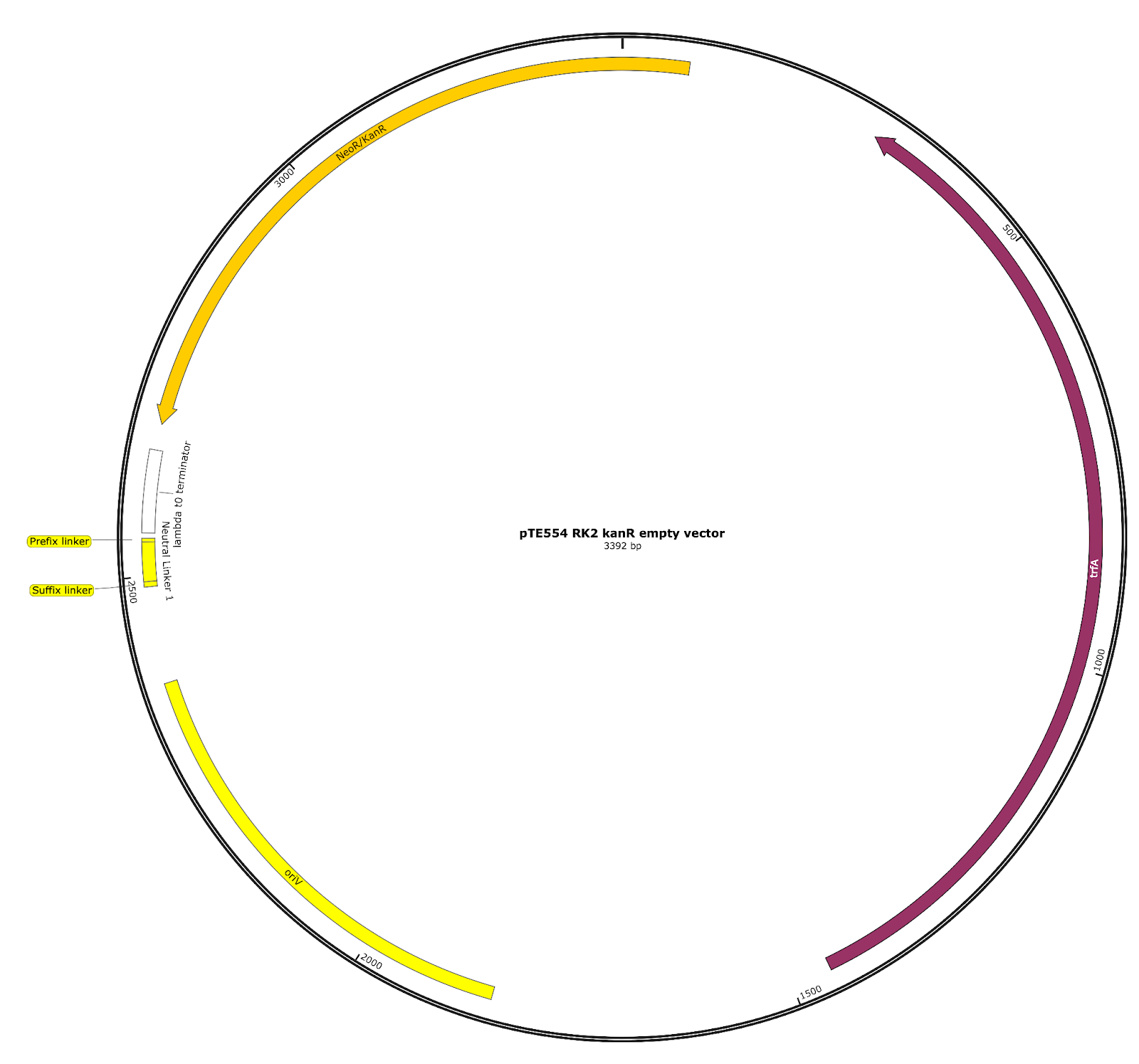

**Supplemental Figure 2.** Plasmid map of the empty vector used to build promoter and RBS libraries containing origin of replication RK2 and kanamycin resistant marker. The plasmid DNA sequence is available on the public JBEI repository.

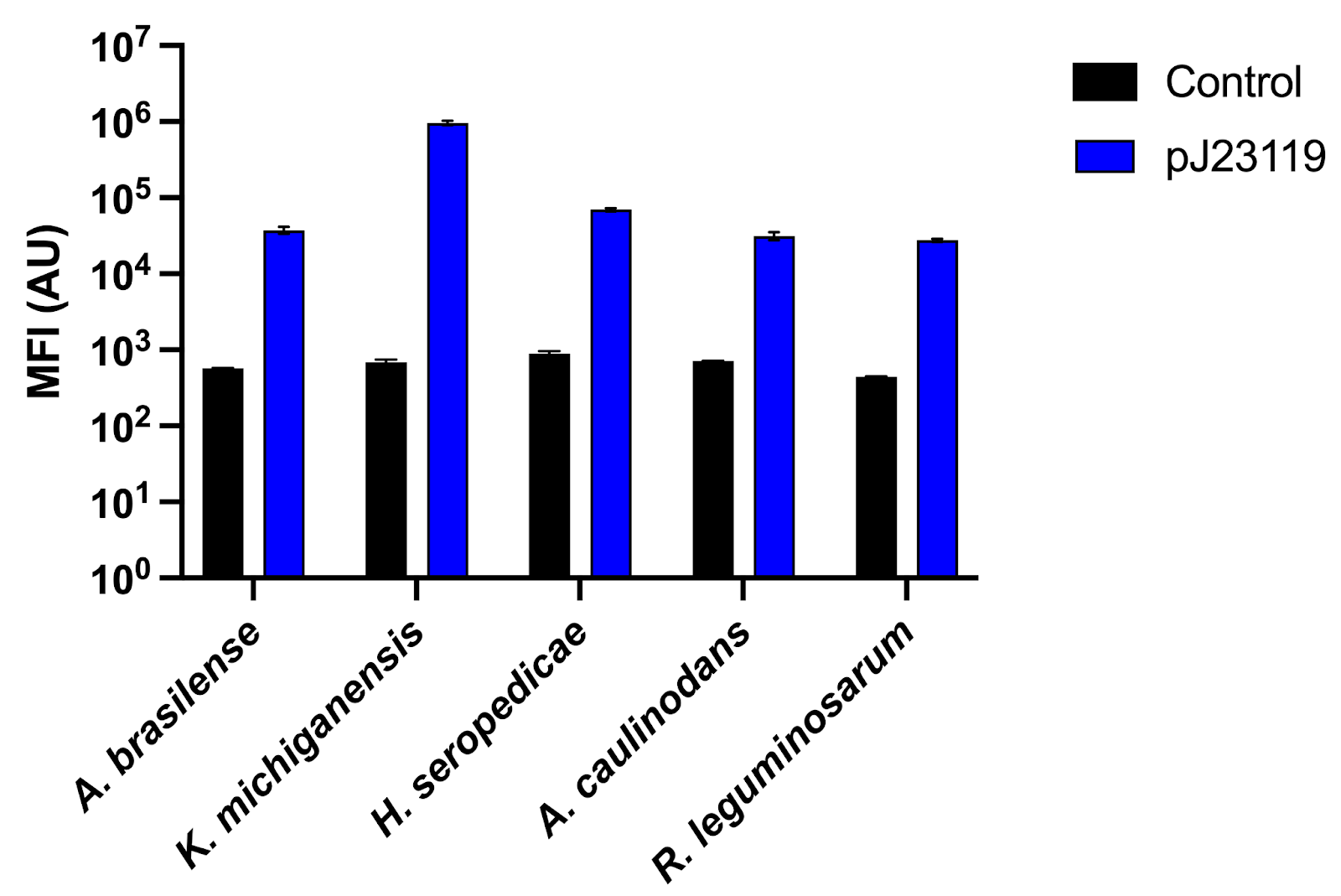

**Supplemental Figure 3. Constitutive expression under Anderson promoter pJ23119.** Expression of GFP under control of the constitutive Anderson promoter J23119 (blue, pTE740) compared to a negative control (black, pTE741).

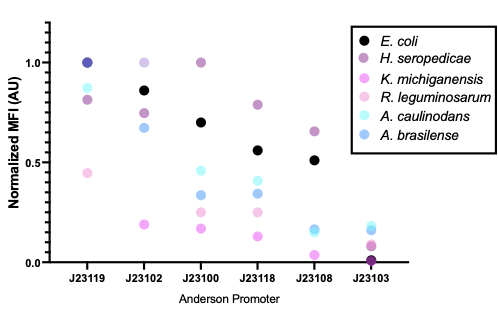

**Supplemental Figure 4. Inconsistent relative expression between promoter sequences in proteobacteria.** Normalized sfGFP fluorescence values (highest fluorescence normalized to a value of 1) between *E. coli* and diazotroph strains.

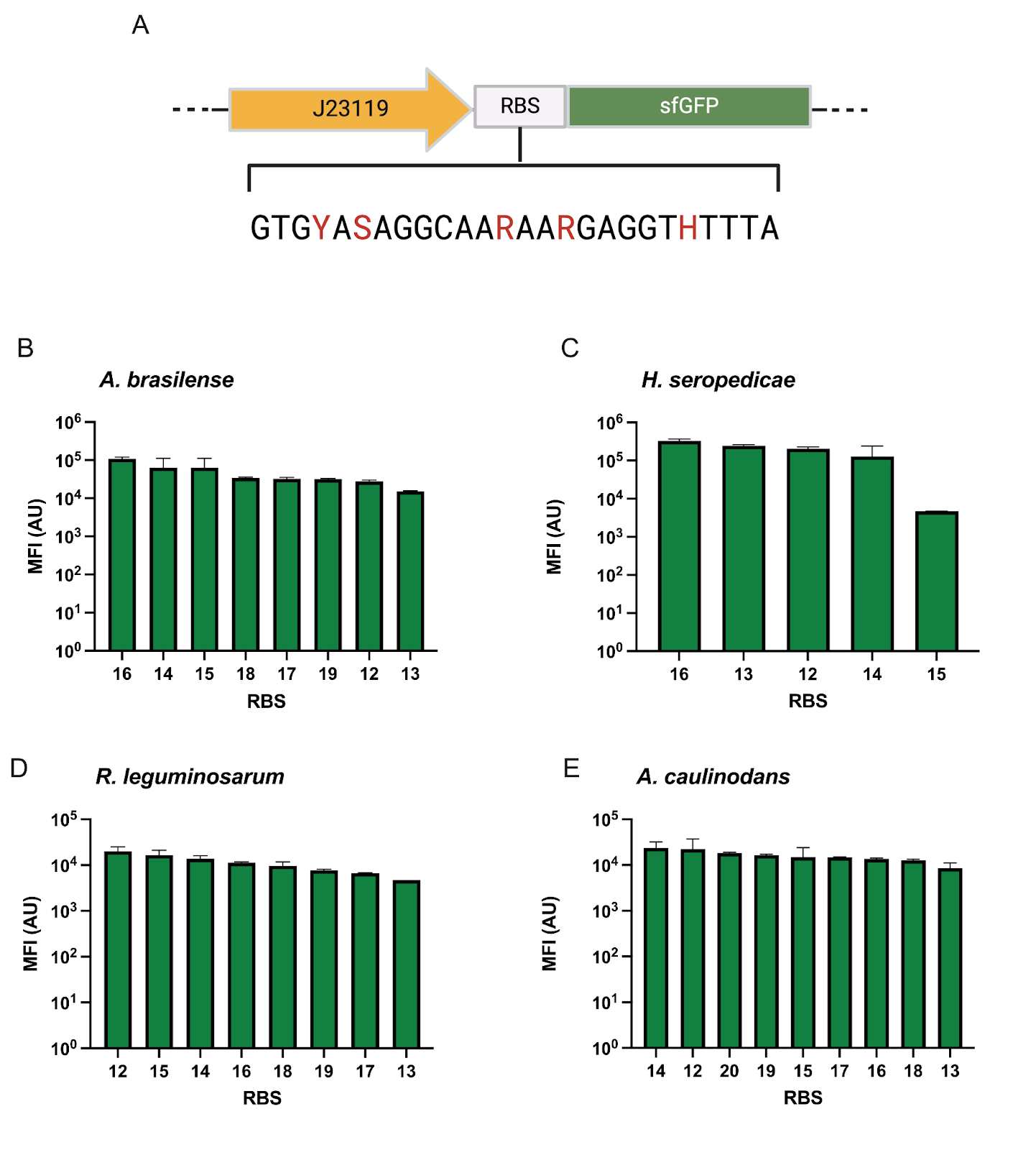

**Supplemental Figure 5. Additional RBS library in four diazotrophs.** Graphical schematic of regulatory components for sfGFP modulation including degenerative oligo used to build the RBS library (A). Mean fluorescence intensity (MFI) of GFP under control of Anderson promoter J23119 and variable RBS (B-E).

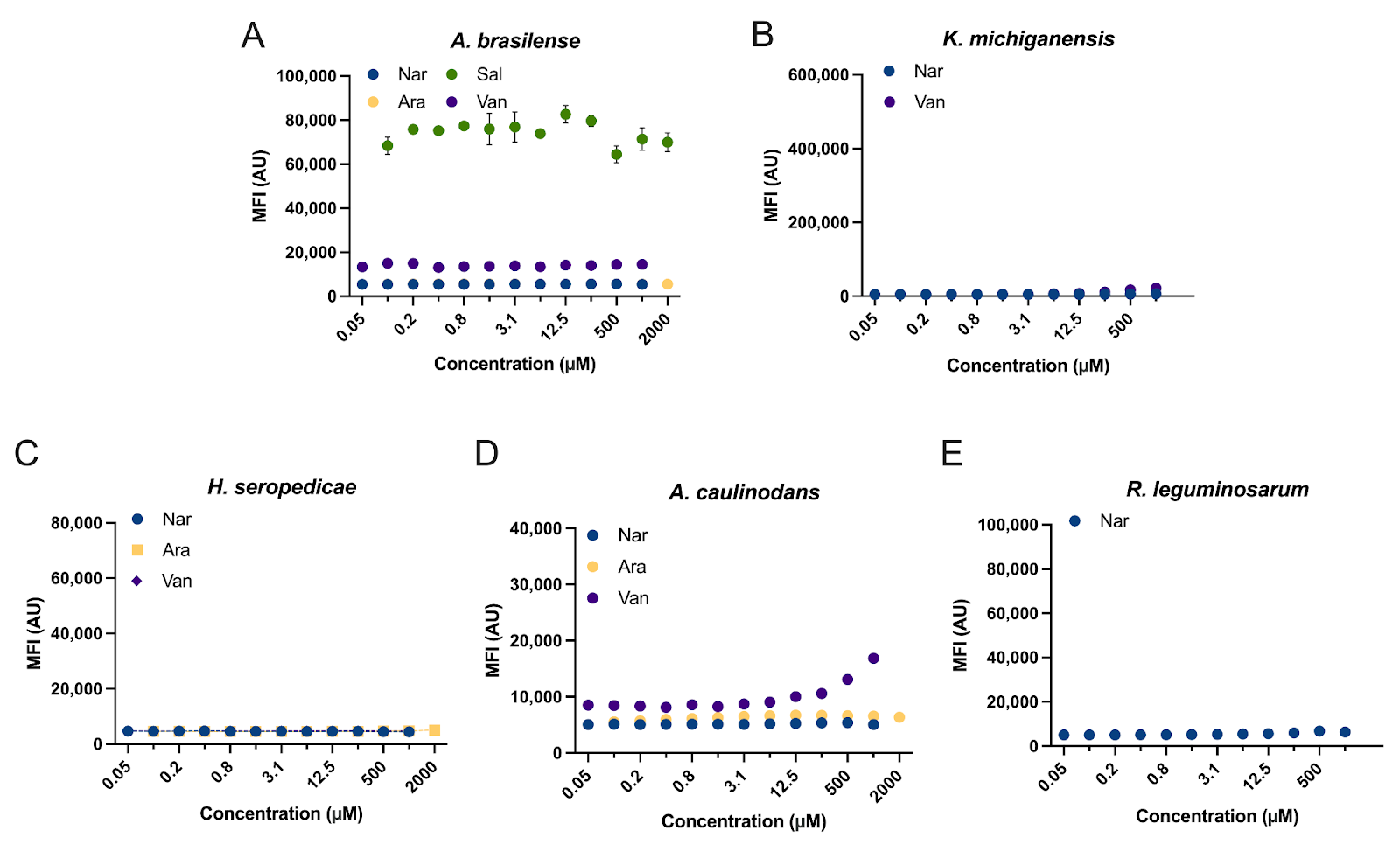

**Supplemental Figure 6. Non-functional inducible systems**. MFI response of different inducible systems shows high background and poor response.

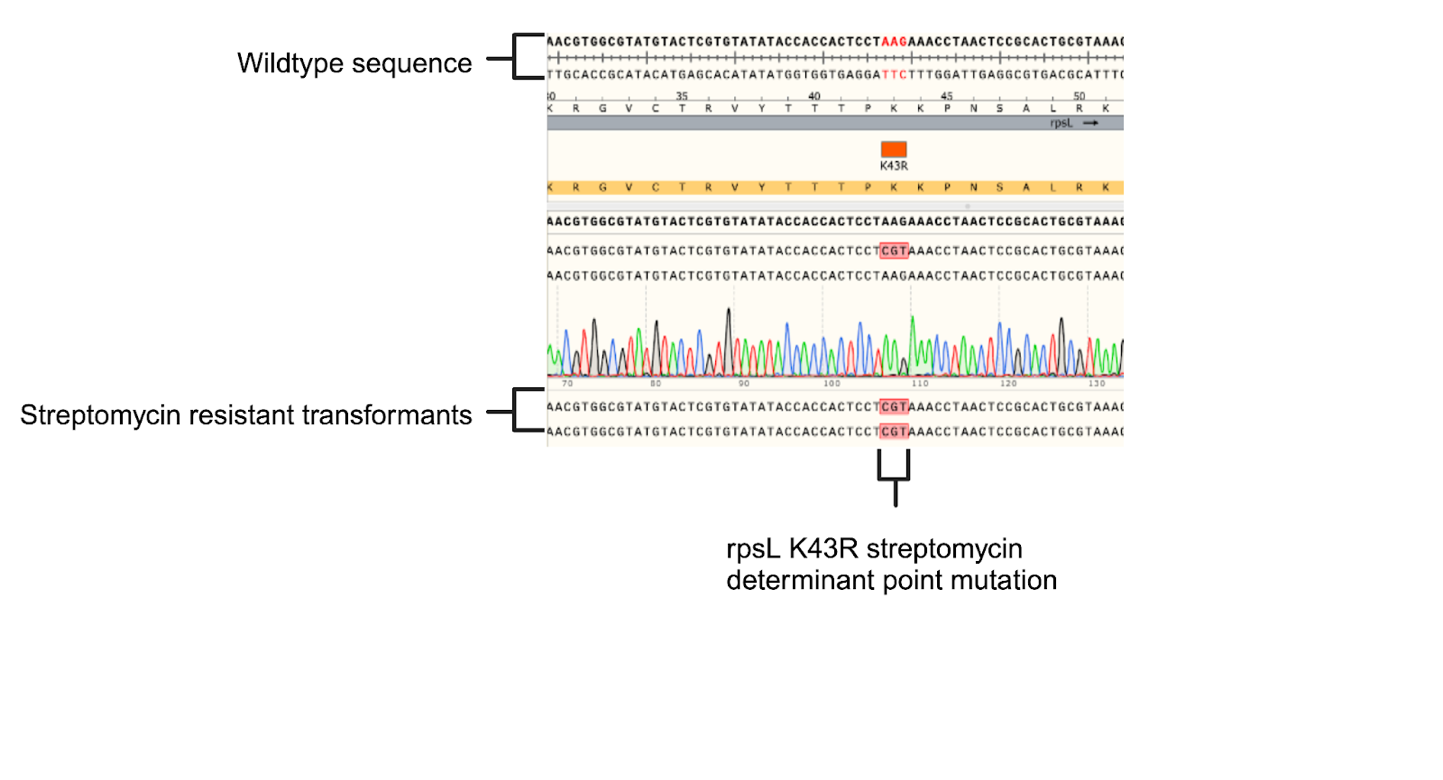

**Supplemental Figure 7.** Sanger sequencing of rpsL K43R point mutation encoded by the mutant codon AAG -> CGT with ssDNA recombineering in *K. michiganensis* m5al.
